# Exogenous Protein as an Environmental Stimuli of Biofilm Formation in Select Bacterial Strains

**DOI:** 10.1101/683979

**Authors:** Donna Ye, Lekha Bapu, Mariane Mota Cavalcante, Jesse Kato, Maggie Lauria Sneideman, Kim Scribner, Thomas Loch, Terence L. Marsh

## Abstract

A screening of environmental conditions that would elicit robust biofilm in a collection of *Serratia marcescens* isolated from soil revealed that exogenous milk protein increased biofilm productivity up to ten-fold. A select screening of fish pathogens, freshwater and human isolates identified several other species that responded similarly to exogenous protein. The optimal protein concentration was species specific; *S. marcescens* at 5% milk protein, *Aeromonas* sp. at 2-3%, *Flavobacterium columnare* at 1% and *Pseudomonas aeruginosa* at 0.1-0.4%. Media supplemented with milk protein also increased the cell counts in biofilm as well as the protein incorporated into the biofilm matrix. These data suggest that relatively high concentrations of exogenous protein may serve as an environmental trigger for biofilm formation, particularly for pathogenic bacteria exposed to relatively high concentrations of protein in bodily fluids and mucosal surfaces.

## INTRODUCTION

Since the concept of biofilm was first viewed through the eyes of molecular microbiology three decades ago, our appreciation of its importance in ecology has grown exponentially. We now recognize biofilm as an alternative life strategy for many, if not all species of microbes across three domains. The importance of biofilm in industry(1), public health (2), medicine(3–5) and the environment(6, 7) have been well documented, leading us to frequently include the analysis of a bacterial strain’s ability to form biofilm as part of its species’ description.

Growth of a microbe in a biofilm removes it from a pelagic lifestyle that is characterized by mass action events (or close to it). As a pelagic entity, change in location and concomitant change in access to nutrients could happen quickly. Once attached to a surface and embedded in a macromolecular matrix, a very different lifestyle ensues – one in which the tempo and mode of life is slowed, the ambient conditions change more gradually (in general) and cell physiology is altered (8).

Biofilm, as part of a life history, is reversible (3, 9, 10). A pelagic microbial species can attach to a substratum, multiply and then return to the surrounding solution as the biofilm structure becomes more susceptible to turbulence or the bacterium senses suboptimal conditions for sessile life. Environmental signals play an essential role in informing the microbe so that optimal survival strategies are selected for, including attachment and release. Consistent with the substantial phylogenetic and physiological diversity of the microbial world, biofilm, as an eco/evo strategy, has been employed in many different ecosystems in response to a broad range of environmental conditions. Of interest to us are specific environmental signals that elicit behavioral changes leading to the formation of biofilm. There are logical candidates for signals including carbon and energy sources, essential micronutrients and even inhibitors. For example, in *Janthinobacterium*, both violacein and biofilm production were stimulated by glycerol and inhibited by glucose (11). The presence of calcium ion has also been found to influence biofilm productivity in *Pseudomonas* (12), *Pseudoalteromonas* (13), *Xyella* (14) and *Citrobacter* (15). In addition, CaCl_2_, MgCl_2_, CuSO_4_, sucrose and sodium dodecyl sulfate produced greater biofilm in *Yersinia pestis* (16). In some circumstances it is difficult to distinguish between a primary trigger and a secondary adjuvant of biofilm formation.

We report herein on two converging lines of investigation in our laboratory, the identification of environmental signals for biofilm formation in a collection of *Serratia* isolated from soil, and in several fish bacterial pathogens. Our observations with *Serratia* clearly indicated that exogenous skim milk protein at relatively high concentrations was sufficient to stimulate biofilm formation ten-fold. We extended this observation to *Flavobacterium columnare*, *Aeromonas salmonicida*, two fish pathogens, as well as a collection of freshwater isolates. We also observed that exogenous protein had little or no effect on some isolates from mammalian hosts, but stimulated others. These data indicate that exogenous protein promote biofilm production in select strains of bacteria.

## MATERIAL AND METHODS

### Strains and cultivation

*Pseudomonas aeruginosa* PA01 was a gift of Dr. M. Bagdasarian (MSU). *Hydrogenophaga F14*, *Brevundimonas F16*, *Acidovorax* F19 and *Pseudomonas* strain C22 were isolated from lake sturgeon eggs (17) and *Serratia marcescens* strains RL-1-RL16 were selected on Pseudomonas Isolation Agar (Difco) from soil under an arborvitae on the Michigan State University campus in East Lansing, MI (GenBank Accession MF581042-MF581057). The strains described in Figures 2 & 3 were isolated from the Red Cedar River on the Michigan State University (MSU) campus by selection on Pseudomonas Isolation agar (GenBank Accession MG386765-MG386811). *Flavobacterium columnare* 090702-1 and *Aeromonas* strain 060628-1 were provided by Dr. Thomas Loch at MSU and were from fish necropsies. *Aeromonas* strain SM, unpigmented *Serratia* and the low biofilm forming *Escherichia coli* were isolated from human feces (18) and generously provided by Dr. Shannon Manning at MSU, as was the high biofilm forming bovine *E. coli* isolate. Isolates were stored at −80°C and resuscitated on Trypticase Soy Agar (Difco) or R2A (Difco). Broth cultures when needed were grown on either TSB or R2B (Difco R2A recipe without agar). Media supplemented with milk protein (Hardy Diagnostics) was made by first sterilizing 2x media stocks (TSB or R2B) and 2x milk protein in separate bottles and then mixing the two shortly after removing the liquids from the autoclave.

### Phylogenetic analysis

The rRNA of *S. marcescens* strains (RL-1-RL-16) specific to this study were sequenced at the MSU genomics facility using the 27F 16S rRNA primer (Sanger chemistry). The *Serratia* sequences (RL-1 through RL-16) were initially screened with the Ribosomal Database Project *Classifier* and *Sequence Match* (19). Phylogenetically related Proteobacteria sequences were downloaded from the Ribosomal Database project and analyzed along with the *Serratia* sequences (RL-1 – RL-16) in SeaView V4 (20) using BioNJ with HKY distance correction and Maximum Likelihood. The maximum likelihood tree and results from *Classifier* and *Sequence Match* are presented in the supplementary materials. All strains isolated from the Red Cedar River were identified by 16S rRNA sequencing using the 27F primer and identified using *Sequence Match* and *Classifier* in the Ribosomal Database Project (19).

### Standard Biofilm Assay

The protocol for biofilm measurement as described by Merritt et al. (21) was used with the following modifications. We used either 96 well or 24 well microtiter plates (Corning Costar) depending on the specific experimental requirements. The 24 well plates were used in experiments when growth, cell and protein concentrations of biofilms were to be determined. Overnight cultures of strains in either Typticase Soy Broth (Difco) or R2B (3-5 mls) were grown at 25°C in a rotating rack (Cole-Parmer). Sterile broth (75-100µl) was added to all wells and then 50-75µl of broth culture was inoculated into the wells. In all experiments, the amount of culture and broth totaled 150 µl for 96 well plates. When using 24 well plates for cell growth measurements, 750µl of sterile broth and 50 µl of overnight culture were added to all wells, excluding the uninoculated controls. When using 24 well plates for cell and protein concentrations, 700 µl of sterile broth and 100 µl of overnight broth culture were added to all wells, excluding the uninoculated controls. After inoculation, plates were sealed with sterile foil (VWR) and incubated at 25°C on an orbital shaker (100 rpm) for 24 or 48 hours depending on the experiment. After incubation, the seal was removed and when processed for biofilm determination, the plates were washed gently (x3) in reverse osmosis (RO) water as described by Merritt et al. (21), stained with 150 µl (800 µl for 24 well plates) of 0.5% filtered (0.2 µ filter) crystal violet for 15 minutes, washed x3 in RO water, blotted and allowed to dry overnight in the dark. The following day 150 µl of 30% acetic acid was added to each well (800 µl for 24 well plates) and the plate was incubated for 25 minutes at 25°C shaking at 100 rpm. Absorbance at 595 nm was measured in a Biotek EPOCH plate reader with 2 measurements for each well. Each sample had at least four replicates within the plate and each media formulation had at least four uninoculated wells that served as negative controls. The average absorbance of uninoculated wells was subtracted from sample biofilm wells.

### Measuring cell growth in milk protein supplemented media

To test for the effects of milk protein on growth of *P. aeruginosa, S. marcescens, Aeromonas sp.* and *F. columnare*, we measured growth in R2B supplemented with 0.1%, 0.2% and 0.4% milk protein in 24 well microtiter plates. We intentionally selected low concentrations at which milk protein is completely soluble. At higher concentrations (>1%) milk protein solutions are colloidal, making it difficult to measure optical density. Growths were performed in 24 well microtiter plates with 4x replication and shaking at 100 rpm on an orbital shaker at 25°C. Optical density measurements were made at 0, 110 min, 210 min, 300 min 390 min 450 min and 24 hours on a Biotek EPOCH plate reader at 600nm. At 24 hours the wells were tested for biofilm formation as described above. Uninoculated controls were subtracted from growths at each time point and in biofilm quantitation. Uninoculated controls for each protein concentration were also replicated x4.

### Determining Viable Cells Within Biofilm

To determine the viable cell count within biofilm formed in supplemented and unsupplemented media, cultures of *P. aeruginosa* PA01, *Aeromonas* 060628-1 and *S. marcescens* RL-5 were established in 24 well plates by inoculating 700 µl of R2B ± milk protein with 100 µl of overnight culture. Control wells contained 800 µl of uninoculated media. Plates were sealed with sterile foil and incubated for 24 hours shaking at 100 rpm and 25 °C. These experiments were set up in duplicate so that biofilm determination with crystal violet (Sigma) and viable cell counts could be performed in parallel. Each plate contained four replicates of each strain and media combination. After 48 hours of growth, one plate was stained with crystal violet as described above for quantitation of biofilm and the duplicate plate was used to determine the viable cells count within biofilms, as follows. The plate was gently rinsed three times in sterile water and the washed biofilm was scrapped off using 600 µl of sterile water and a sterile applicator. The cell slurry was transferred to 1.5 ml eppendorf tubes, vortexed to break up cell aggregates and 10-fold serially diluted for plating onto R2A. Plating was in triplicate and CFUs are reported as the total CFUs per well.

### Determination of protein content in biofilm

To determine the protein concentration of biofilms, 24 well plates were used as described. In this experiment we tested *P. aeruginosa*, *Aeromonas* 060628-1 and *F. columnare*. Biofilm of *P. aeruginosa* and *Aeromonas* were prepared in R2B and R2B-5%MP while *F. columnare* was tested in R2B-1%MP. Duplicate plates were inoculated so that both protein concentration and crystal violet staining could be tested in parallel. In each plate, all unique media conditions were replicated four times. After 24 hours of incubation at 25°C and shaking at 100 rpm, one plate was stained with crystal violet (as described above) and the duplicate plate was used to determine protein content within biofilm as follows. After 24 hours of incubation the media was removed and the wells were washed twice by adding 800 µl of sterile water and shaking at 100 rpm for 2 minutes. The final wash was removed and 200 µl of sterile water was added to each well. The biofilm was removed by manually scrapping with a sterile glass rod and then transferred to a 1.5 ml Eppendorf tube. The solution was vortexed and centrifuged at 4°C and12,000 rpm for 20 minutes in a microfuge to remove the cells. The supernatant was transferred to a new tube and 400 µl of 100% Ethanol was added to precipitate protein. After overnight storage at −20°C, the tubes were centrifuged at 4°C and 12,000 rpm for 20 minutes to pellet all protein. The supernatant was decanted and the pellets were air dried for 15 minutes and then resuspended in 150 µl of 1x PBS. To determine the protein concentration in these samples the Coomassie protein assay (Thermo-Scientific) was employed, using the vendors recommended protocol. Briefly, 150 µl of Coomassie reagent plus 150 µl of sample was added to a microtiter plate well, mixed and incubated for 10 minutes in the dark. The plate was read at 595 nm using a Biotek EPOCH plate reader. The standard curve was as recommended by the vendor.

### Confocal microscopy

A two-well chamber (Lab-TekII, Nalge Nunc International, USA) was inoculated with 700 μL of media and 100 µl of *P. aeruginosa* or *F. columnare*. 5% Skim Milk Protein at 1/2x Tryptic Soy Broth (Becton, Dickinson and company, France) medium and 100 μL of *F. columnare* overnight broth. The chamber was then wrapped in Parafilm (Bemis, USA) to seal it. Next, the sample was incubated for 72 hours at 100 RPM. At 48-hour incubation the media was gently removed and new 700 μL of fresh medium was added into the wells. At 72 hours, the medium was removed and discarded, and the chamber was gently washed 3 times with 1 mL of sterile water. 1 mL of fluorescent solution containing 0.5mL of 20x Nano Orange dye (Molecular Probes Protein Quantitation Kit N10271) and 0.5mL of FM4-64 dye (Molecular Probes FM4-64) was added into one well and incubated in the dark for five minutes. The well was washed with 1 mL of sterile water two times. The sample was kept hydrate during microscopy. A confocal microscope (Olympus FluoView FV1000) was used for imaging at 20x and 90x.

## RESULTS

During the screening of a variety of environmental conditions designed to stimulate the formation of biofilm by soil isolates of *Serratia*, we detected a significant increase in biofilm when our standard growth media, R2B, was supplemented with milk protein (MP). Using standard media-grade skim milk protein (Hardy Diagnostics) at 5%, biofilm formation of 16 independently isolated soil *Serratia* strains increased significantly. In Figure 1 we show the response of five *Serratia* strains to media supplemented with different concentrations of milk protein (0.05, 0.5, 2.5 and 5.0%). All five isolates responded to 5% MP supplementation with a ten-fold increase in biofilm formation. All remaining isolates responded with similar increases (data not shown). In our biofilm assays we routinely use *Pseudomonas aeruginosa* PAO1 as a positive control. Under all conditions we have tested, PA01 produced a robust biofilm when grown on R2B or TSB but had little response to the presence of exogenous protein when at 0.5% or greater and, in many of our assays, high concentrations of protein in the media slightly inhibited biofilm formation by *P. aeruginosa*.

**Figure 1.**
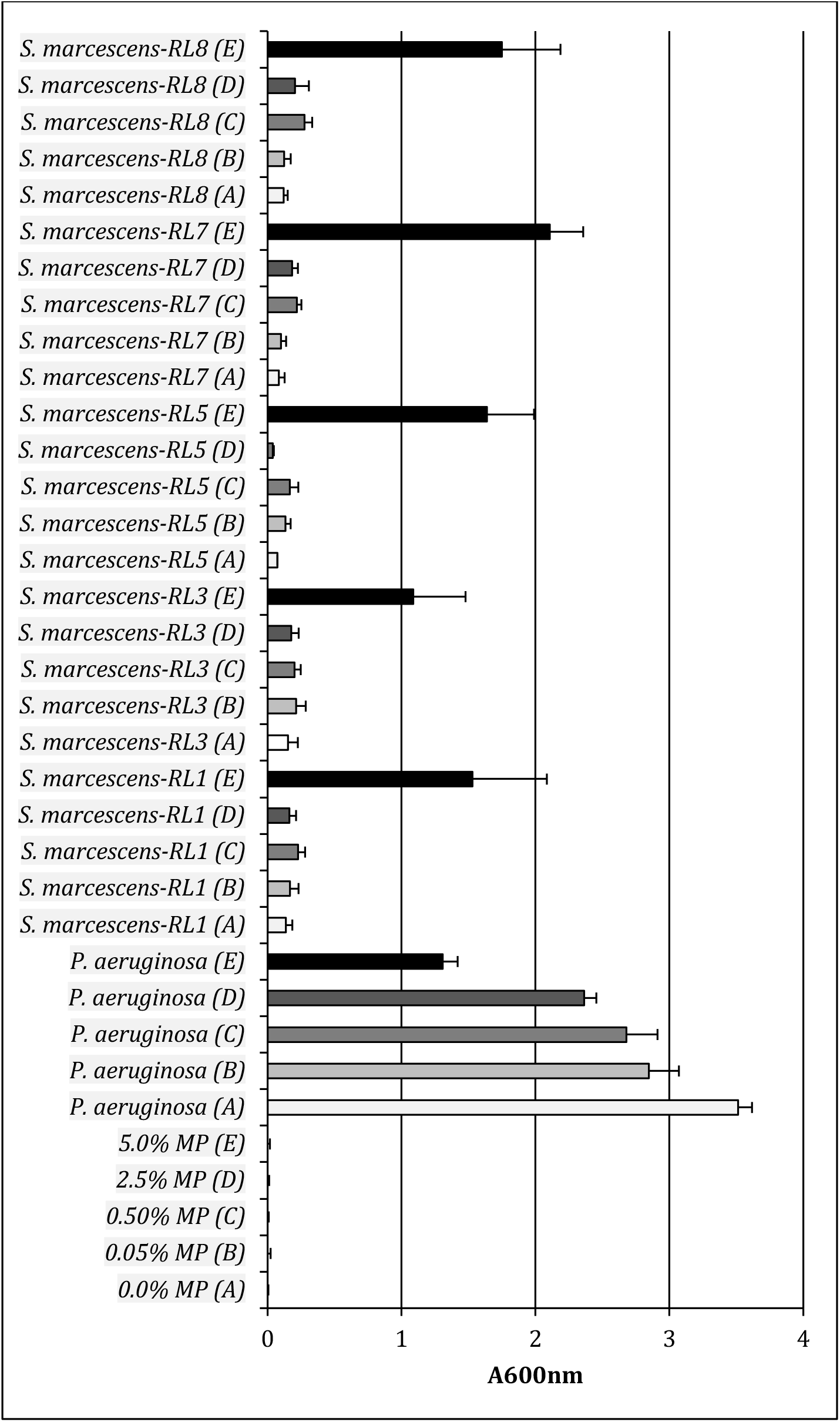
Biofilm formation by *S. marcescens* soil isolates in response to elevated concentrations of milk protein in broth (0.05%, 0.5%, 2.5% & 5.0%). Incubation in microtiter plates was for 24 hours at 25°C on an orbital shaker at 100 RPM.

To extend these observations to other species we tested the ability of 5% MP to stimulate biofilm production in 200 freshwater bacterial isolates. In Figure 2 we report on 48 randomly picked isolates that were selectively isolated on *Pseudomonas* isolation agar and were therefore resistant to irgasan, a broad spectrum antimicrobial that targets fatty acid synthesis in bacteria. Among these isolates, 14 showed substantial increase in biofilm production in milk protein supplemented media at least 2-fold greater than the unsupplemented control (eg. one *Yersinia*, *Shewanella* and *Rahnella*). Of 13 *Pseudomonas* isolates, only two showed more than a 2-fold increase in biofilm with protein-supplemented media. Of the eight genera in this test, *Aeromonas* consistently showed a robust response to exogenous protein in the media. Six of the twenty *Aeromonas* isolates had 10-fold increases in biofilm formation and 8/20 had at least a 2-fold increase.

We have tested several hundred freshwater isolates in this manner and when assaying that many strains, we routinely make a single-pass evaluation with the crystal violet assay, accounting for the lack of error bars in Figure 2. To statistically confirm our results, we examined six isolates from this freshwater collection in greater detail, three *Aeromonas*, two *Rahnella* and one *Pseudomonas* at four different concentrations of milk protein in TSB. These data, presented in Figure 3, show that the *Aeromonas* isolates responded to MP concentrations of 1-5% while a freshwater *Pseudomonas* isolate revealed little response until 5% and *Rahnella* was unresponsive.

**Figure 2.**
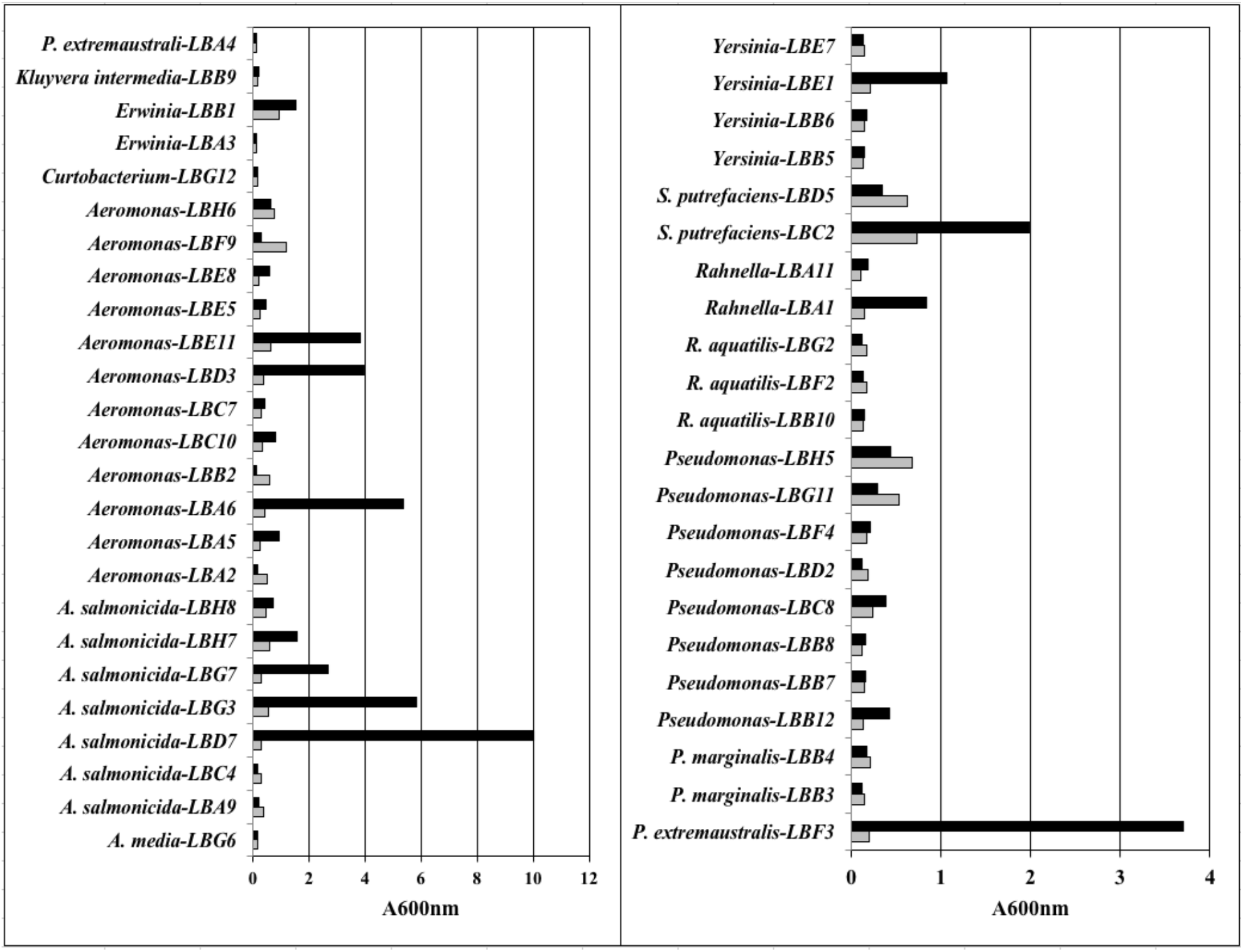
Biofilm formation by freshwater isolates in response to milk protein at 5% in broth. All isolates were from the Red Cedar River, East Lansing, MI. All strains were isolated from a direct plating of river water on Pseudomonas Isolation Agar.

**Figure 3.**
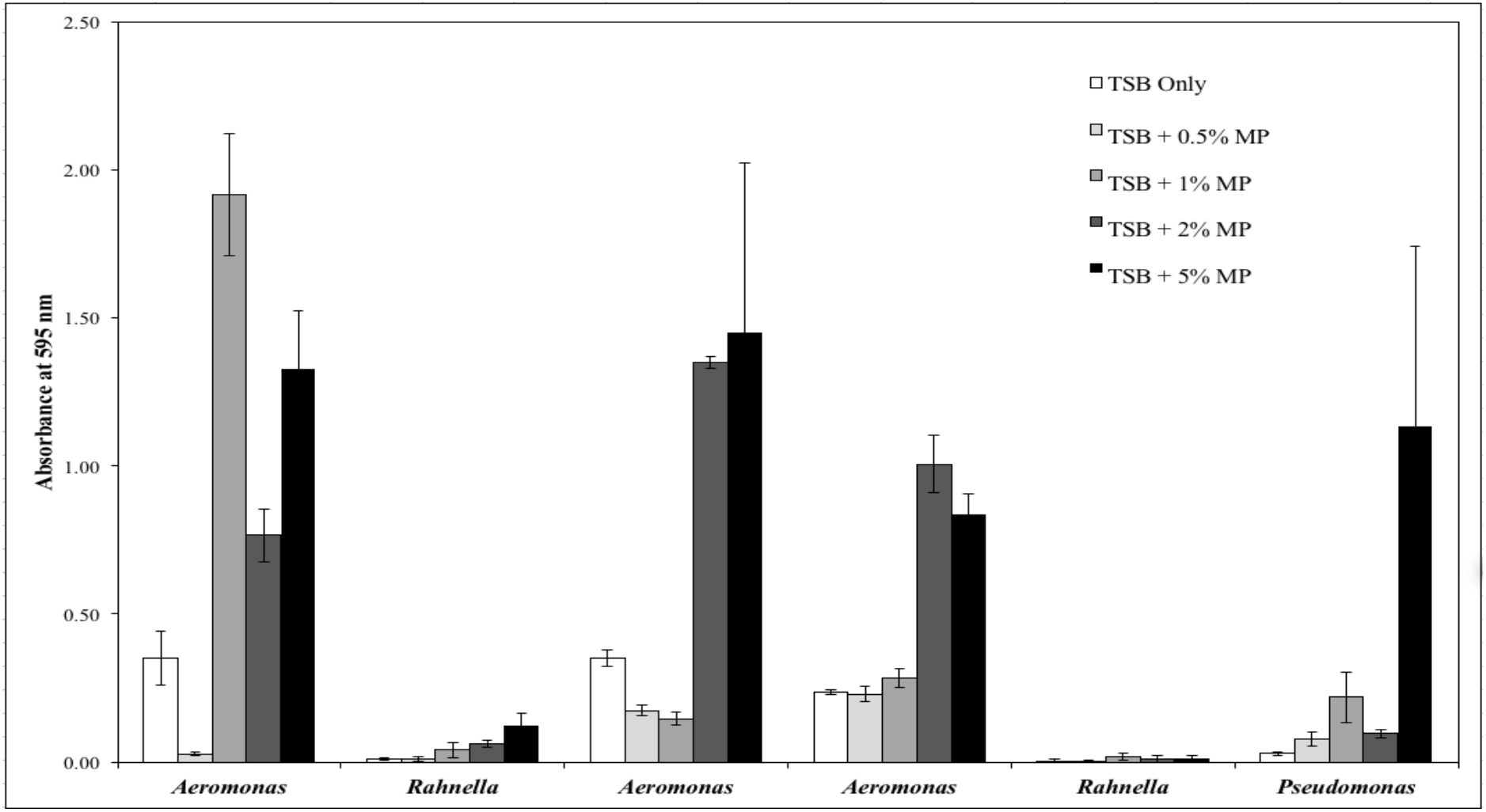
Biofilm formation by six freshwater isolates in response to elevated protein concentrations in broth. Six isolates from the biofilm screening described in Fig.2 were tested for biofilm production at four different concentrations of milk protein (0.5%, 1.0%. 2.0% and 5.0%) in TSB.

To extend this analysis to isolates associated with eukaryotic hosts, we also investigated the relationship between biofilm formation and exogenous protein in five isolates from fish, three from humans and an *E. coli* strain from bovine. These data are presented in Figure 4. Among the isolates from fish, *F. columnare, Hydrogenophaga, Brevundimonas* responded strongly and positively to exogenous protein by producing abundant biofilm, but at different optimal protein concentrations (Fig 4. **Panel A**). *F. columnare* and *Hydrogenophaga* had greatest biofilm productivity at 1% while *Brevundimonas* was more productive at 5%. *Pseudomonas* C22 does not form abundant biofilm in unsupplemented media and productivity increased only modestly at 1% and 5% MP. *Acidovorax* was unresponsive to exogenous protein. **Panel B** reports on the human and bovine isolates. *Aeromonas* sp, an unpigmented *Serratia* and a low biofilm forming *E. coli* were isolated from human feces and had varied response to exogenous protein. Both *Aeromonas* and the *Serratia* isolates responded with greater biofilm productivity at 1% and 5% but *E. coli* was unresponsive. Included in this experiment was one of our soil *Serratia* isolates (RL-4) for comparison. Interestingly, biofilm formation by the *E. coli* isolate from bovine, identified as a high biofilm forming strain, was inhibited in media supplemented with 1% and 5% MP. Note that this experiment was conducted in LB broth without salt to mimic the conditions used in the initial characterizations of the *E. coli* strains. Both *P. aeruginosa* PA01 and our soil *Serratia* RL-4 had biofilm profiles in supplemented and unsupplemented TSB similar to what we have seen in R2B.

As can be seen from these biofilm assays, in some cases the amount of crystal violet staining material was quite large. In many of the *Aeromonas* strains tested an opaque disk formed at the bottom of the wells, particularly if the incubation period was extended to 48 hours and the 96 well format was used. An obvious concern was the possibility that crystal violet was staining protein and biofilm matrix atypically and providing a false positive for biofilm formation. To test for this, we ran several analyses in 24-well microtiter plates that prevented the formation of any opaque disk by virtue of the large well diameter. In these experiments, we measured biofilm formation using crystal violet and performed viable plate counts on biofilm from replica plates. These data are presented in Figure 5 and show that, as observed above, both *Aeromonas* and *Serratia* responded strongly to exogenous protein, producing at least a 10-fold increase in crystal violet signal while *Pseudomonas* had little response. The cell viability from a replica plate revealed a 1.2-3 order of magnitude increase for *Serratia* and *Aeromonas* when grown with 5% exogenous protein, while *P. aeruginosa* had a robust viable count in the absence of protein and only a modest increase with protein when compared with *Aeromonas*.

**Figure 4.**
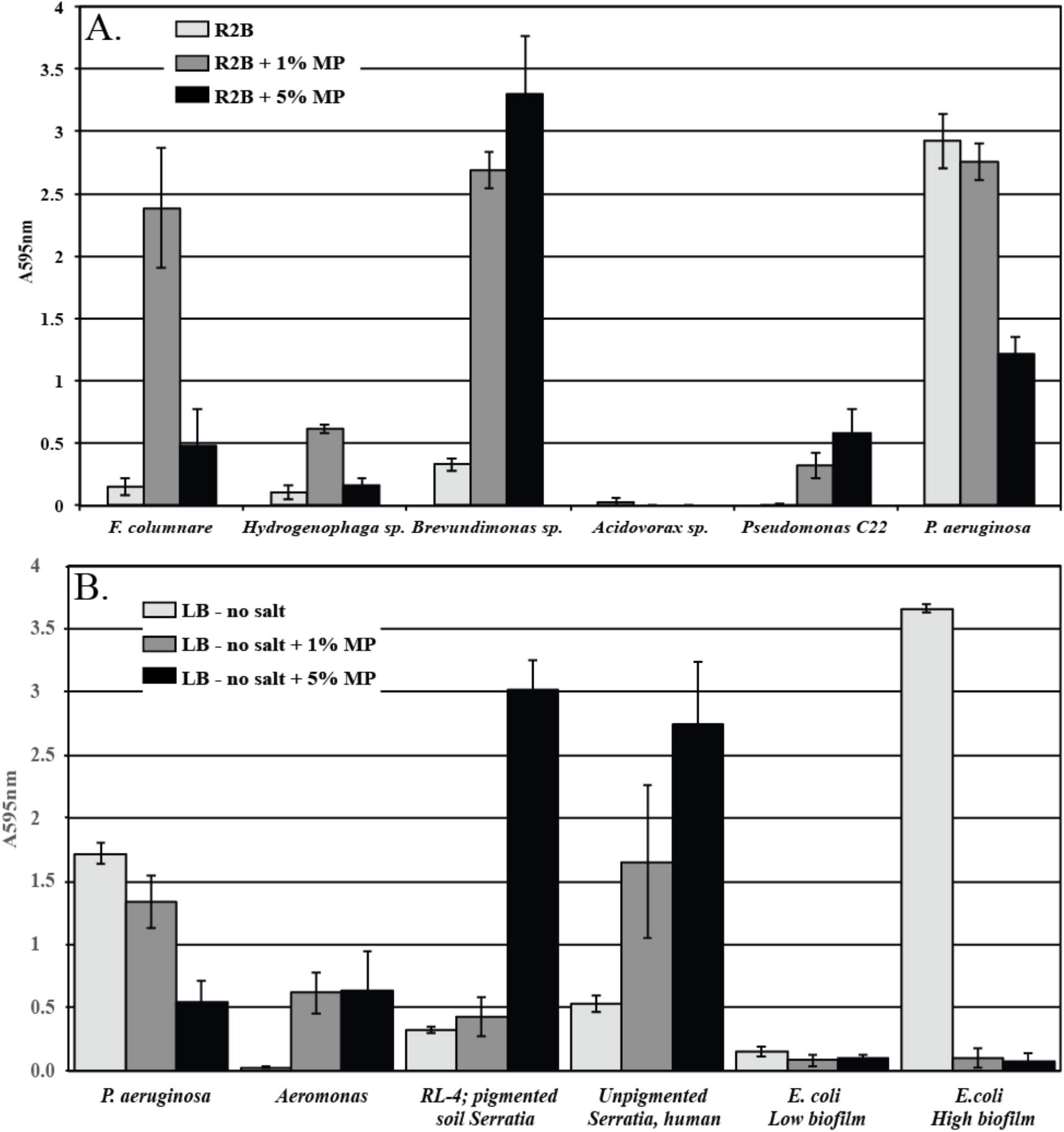
Biofilm formation by bacterial isolates from lake sturgeon eggs, *Homo sapiens* and Bovine. *P. aeruginosa* PA01 was used as a positive control for biofilm formation. Panel A; isolates from fish (*F. columnare* 090702-1*, Hydrogenophaga* F14, *Brevundimonas* F16*, Acidovorax* F19, and *Pseudomonas* C22) tested on R2Broth with 1.0% and 5.0% milk protein supplemented media. Panel B. Biofilm assay performed in LB broth without NaCl at 1% and 5% milk protein on *Aeromonas*, unpigmented *Serratia* and *E. coli* from *H. sapiens* and *E. coli* from Bovine.

**Figure 5.**
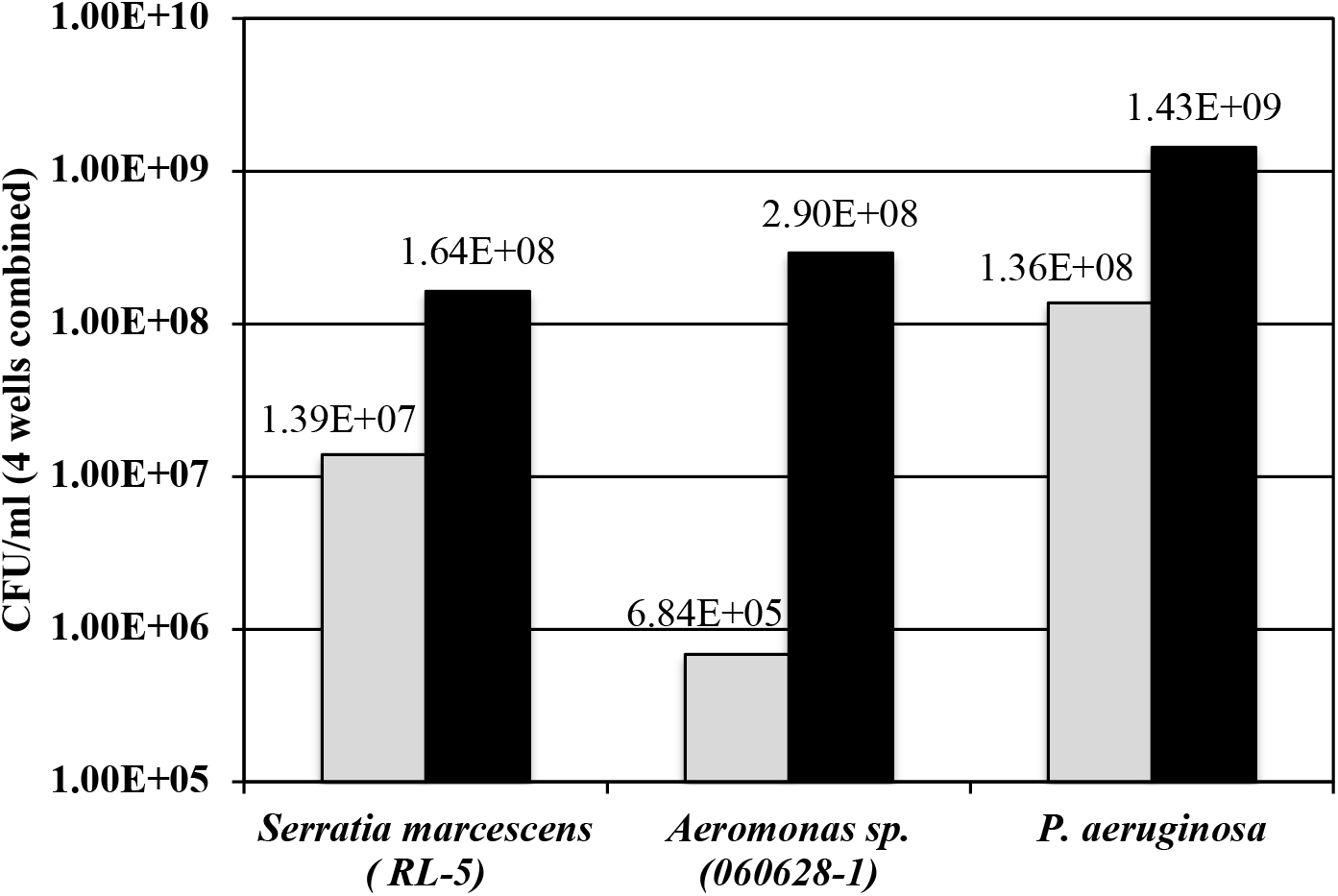
The effect of exogenous protein on the concentration of cells within biofilm. These assays were performed in 24 well plates in R2Broth (gray) and R2Broth supplemented with 5% milk protein (black).

An obvious question regarding the effect of exogenous protein on the formation of biofilm is whether the biofilm becomes enriched in protein. To test for this, we established biofilm in 24 well plates (4 replicates of each strain on a plate) and replicated the whole plate so that both biofilm formation and the amount of protein within the biofilm matrix could be measured. Because of our interest in fish pathogens we tested *Aeromonas* and *F. columnare* with *P. aeruginosa* as our positive control. The results are presented in Figure 6. As shown previously, both *Aeromonas* and *F. columnare* responded strongly to exogenous protein by producing more biofilm while *P. aeruginosa* PA01 was unresponsive. In this test, we used the optimal protein concentrations of 5% for *Aeromonas* (and *Pseudomonas*) and 1% for *F. columnare*. The biofilm from the replica plate was washed and manually scrapped from the wells and the protein concentration was determined using the Bradford assay, after removing the cells by centrifugation. The amount of protein detected in the biofilm for *P. aeruginosa* was 8.2 and 5.4 µg/ml for growth without and with protein, respectively. For *Aeromonas*, the increase in biofilm in response to exogenous protein was accompanied by an increase in matrix protein concentration from 0.55 to 47.3 µg/ml. For *F. columnare*, the 20-fold increase in biofilm was accompanied by a nearly 20-fold increase in matrix protein (0.18 > 3.11 µg/ml). The optical densities of the cultures are revealing as well. As expected, the initial OD of cultures in unsupplemented media was relatively low, representing a 1:8 dilution from overnight cultures, but clear evidence of growth was detected after 24-hour incubation. The initial OD of the protein-supplemented wells was dominated by the opacity contributed by the milk protein, ~1.7 for a 5% solution and ~0.5 for a 1% solution. After incubation for 24 hours the OD of the *P. aeruginosa* wells dropped to 0.45, suggesting the presence of protease activity. Presumptive protease activity was also detected in the *F. columnare* wells, evidenced by a drop in OD from 0.5 to 0.16. Interestingly the wells containing *Aeromonas* showed no reduction in OD.

**Figure 6.**
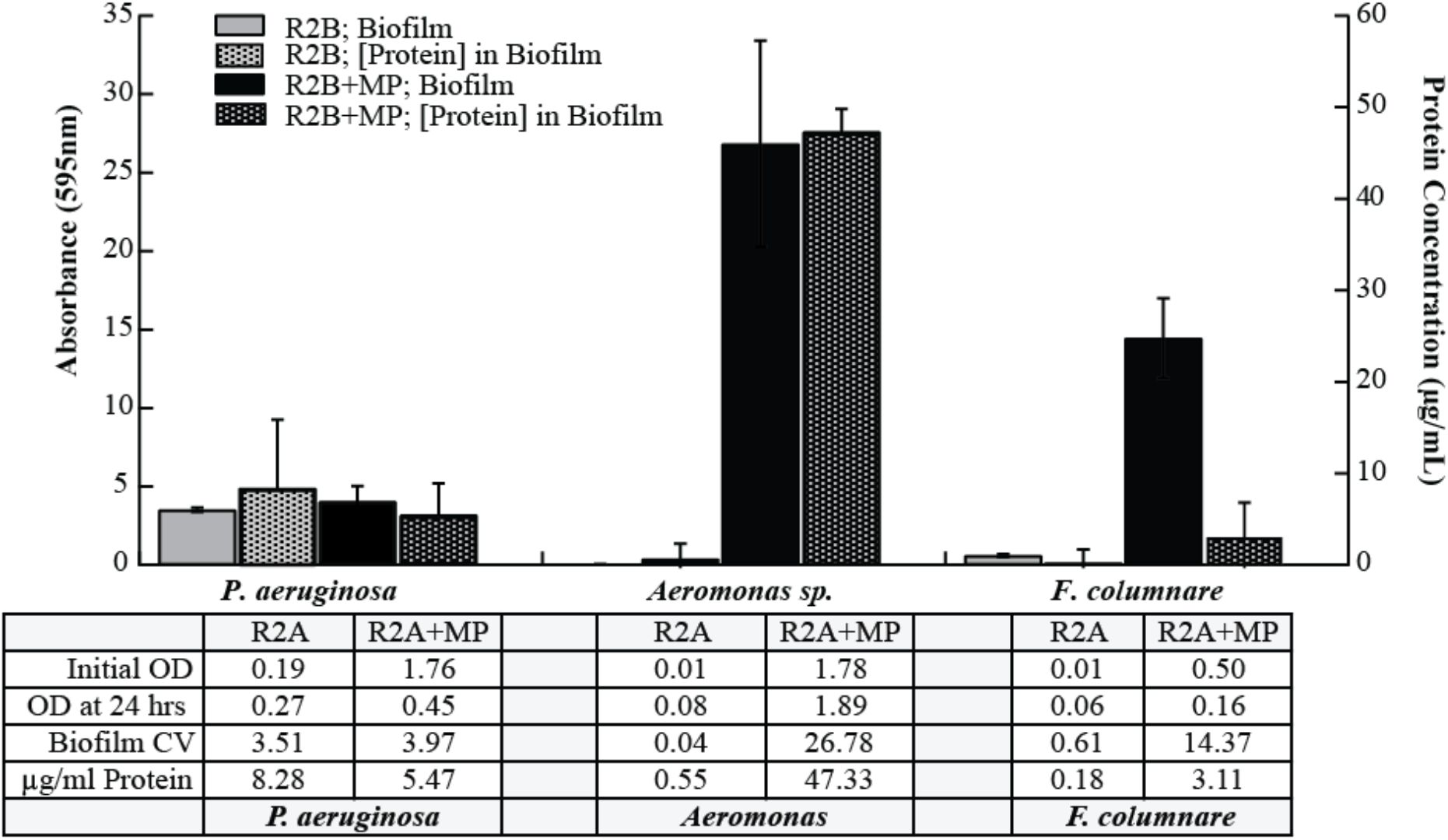
The effect of exogenous protein on the protein concentration within biofilms. These experiments were performed in 24 well plates (4 replicates for each treatment) and each plate was replicated for measuring biofilm (crystal violet) and protein (Bradford assay). *P. aeruginosa* PA01 and *Aeromonas* strain 060628-1 were tested at 5% milk protein and *F. columnare* 090702-1 was tested at 1% milk protein.

To determine the effect of exogenous protein on cell growth we incubated *P. aeruginosa, S. marcescens, Aeromonas* strain 060628-1 and *F. columnare* strain 090702-1 at three concentrations (0.1%, 0.2% and 0.4%) of milk protein in R2B and monitored growth by optical density at 600nm. Low concentrations were selected to avoid colloidal solution conditions present at higher concentrations. *P. aeruginosa* grew well under these experimental conditions but optical density was diminished in a concentration dependent manner when the media was supplemented with protein (**Panel A**, Fig. 7). In contrast, *S. marcescens* (**Panel B**, Fig 7) grew robustly, regardless of the exogenous protein through 450 min. Statistical differences were detected only after 24 hours of growth when exogenous protein appeared to modestly boost growth. *Aeromonas* grew slowly (**Panel C**, Fig 7) through 450 minutes with no appreciable difference with protein addition. The greatest growth was between 450 min and 24 hours. At 24 hours growth was inhibited at 0.4% exogenous protein. *F. columnare* grew well in the absence of exogenous protein and poorly, if at all, in its presence (**Panel D**, Fig 7). The tendency of this strain to form aggregates in solution accounted for the substantial inter-replicate variability. After 24 hours the plates were processed for biofilm formation (Figure 8). *P. aeruginosa* PA01, as mentioned above, is a robust biofilm forming strain. Under conditions of growth in this experiment, enhanced biofilm productivity was detected at all concentrations of exogenous protein. While *S. marcescens* grew vigorously, biofilm productivity was quite low at the tested protein concentrations. *Aeromonas* also lacked biofilm productivity at the lower concentrations of protein but did increase substantially at 0.4% milk protein, in spite of the apparent growth inhibition at this concentration. Biofilm production by *F. columnare* increased in a concentration dependent manner when the media was supplemented with protein. This robust biofilm production was in contrast to pelagic growth which appeared inhibited by exogenous protein.

**Figure 7.**
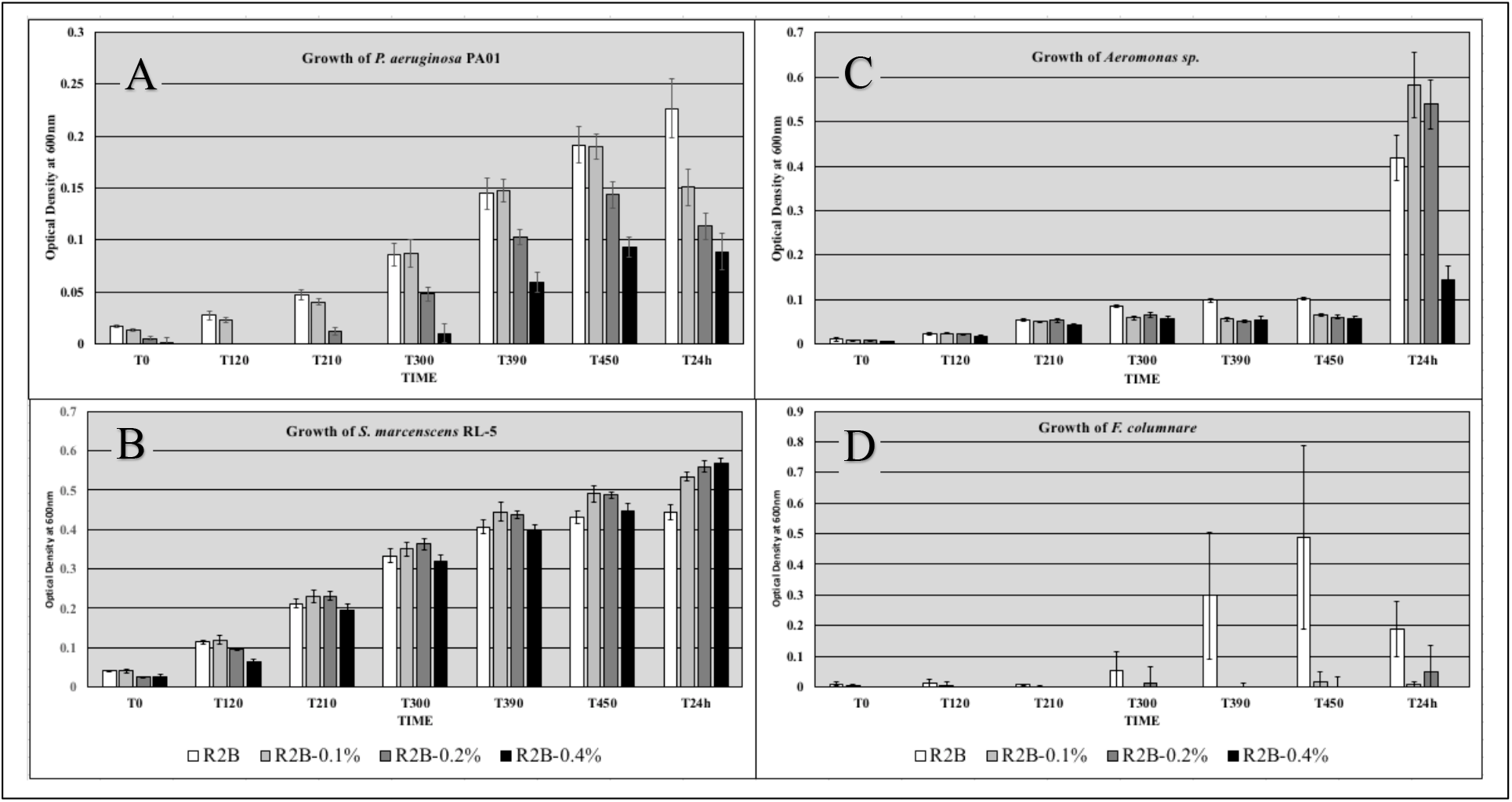
The effect of exogenous protein on growth of *P. aeruginosa* PA01*, S. marcescens* RL-5, *Aeromonas* strain 060628-1and *F. columnare* 090702-1. Growth was measured in R2Broth unsupplemented and supplemented with 0.1%, 0.2% and 0.4% milk protein.

**Figure 8.**
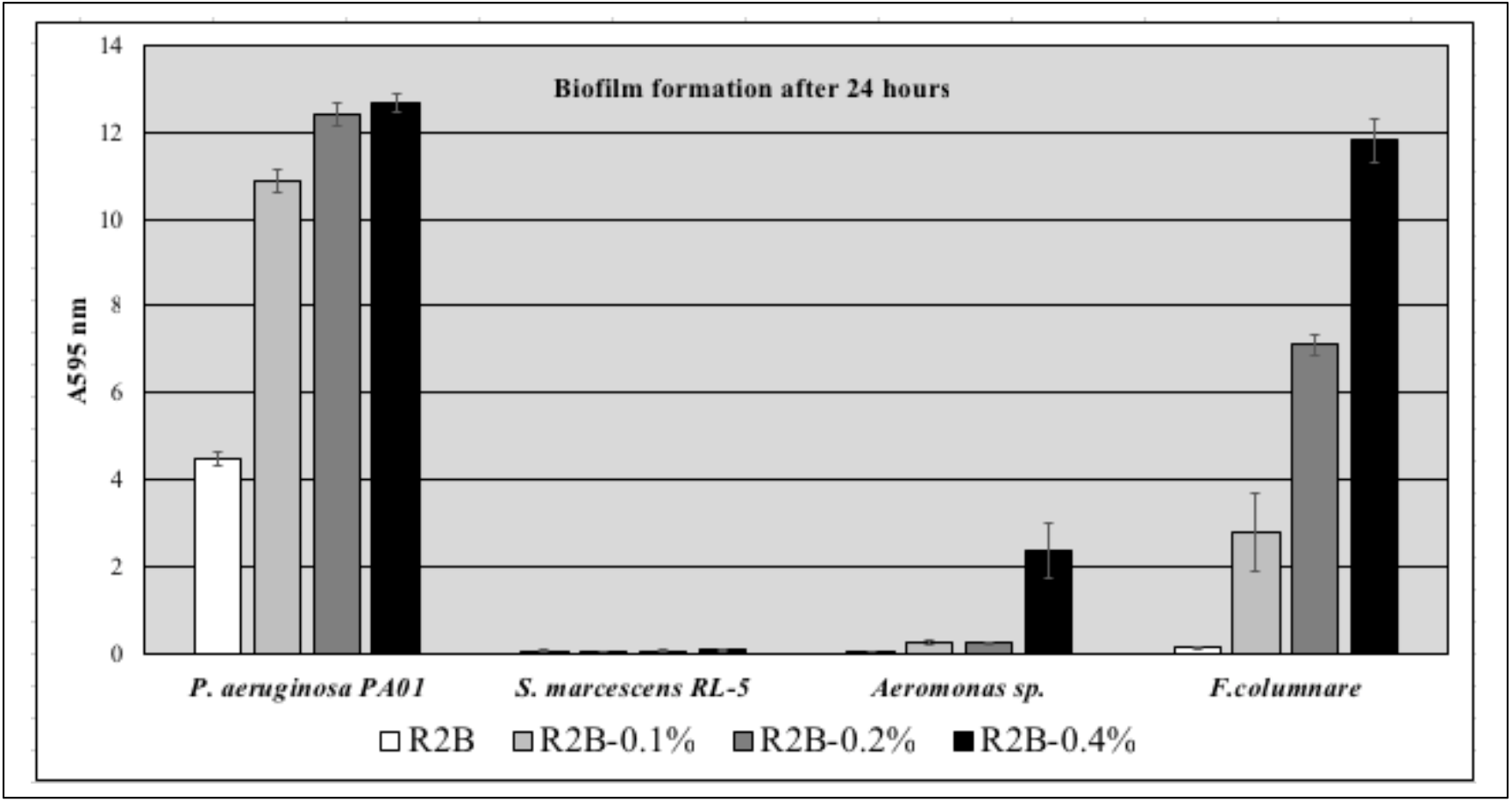
Quantitation of biofilm from growth experiment described in Fig. 7.

In addition to measuring the protein content with a standard Bradford assay we used nano-orange to visualize the biofilm–associated protein. Using the standard microtiter plate protocol, we established biofilm on sterile coverslips with and without exogenous milk protein (1%) using *P. aeruginosa* and *F. columnare* as the test strains. After growth, the biofilm was washed with sterile water (x3) and stained with Nano-orange and FM4-64 using the vendors protocol. The biofilm was viewed on an Olympus FluoView FV1000 Confocal Microscope at 20X and 90X magnification (Figure 9). Numerous *P. aeruginosa* cells were detected at 20X magnification but there was little evidence of a robust contiguous biofilm. Intensely orange spots could be detected suggesting concentrations of protein spotted the surface. At 90X magnification well isolated cells were seen with little evidence of a protein matrix. In contrast, the biofilm formed by *F. columnare* showed a thick branched proteinaceous complex at 20X magnification. Cells were clearly outlined with the lipophilic FM4-64 stain at 90X magnification and showed morphological variation as describe previously (22). In addition, irregularly shaped orange forms as well as cells decorated with Nano-orange were detected.

**Figure 9.**
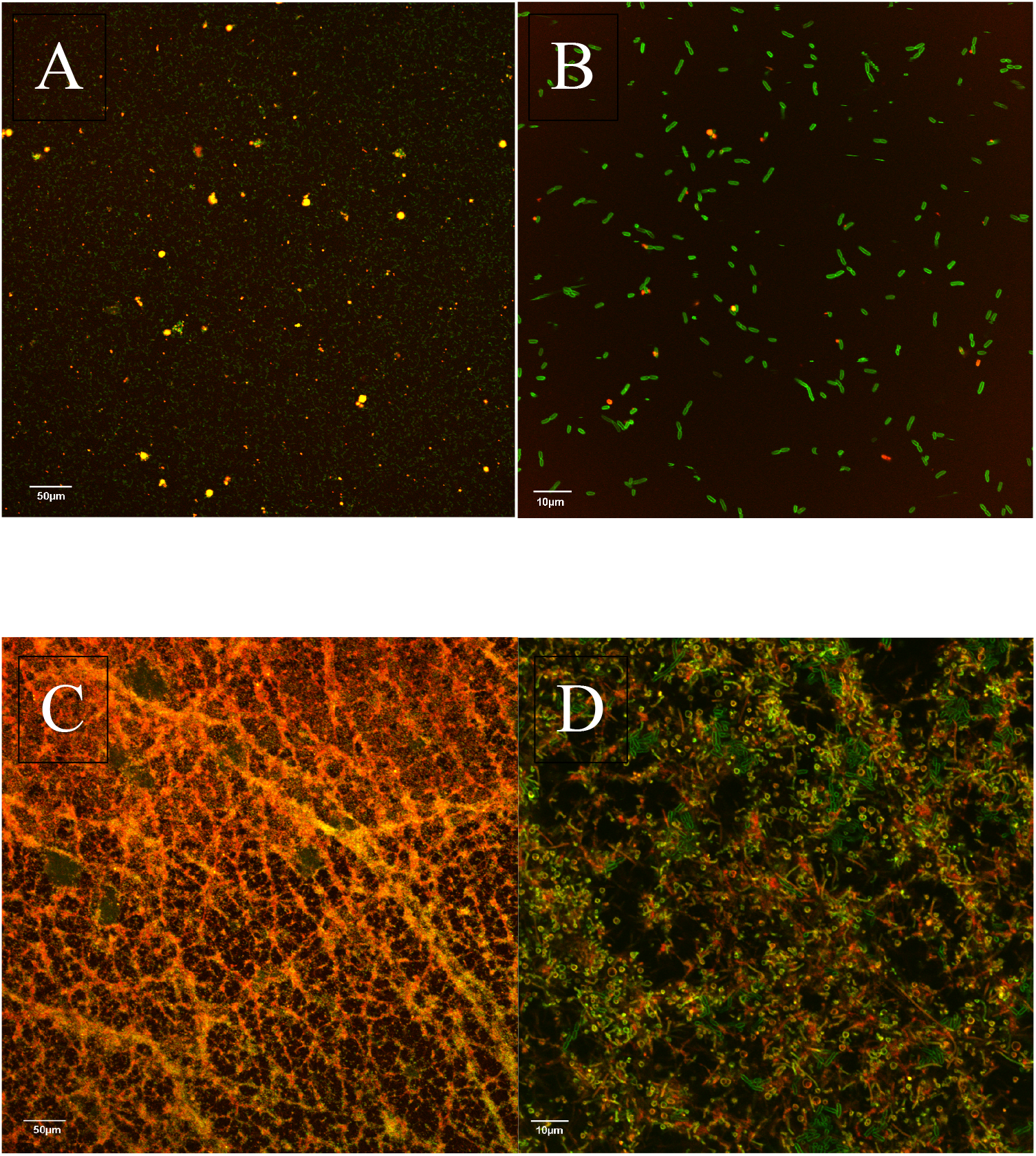
Confocal images of *P. aeruginosa* (A & B) and *F. columnare* (C & D) biofilm at 20X and 90X grown with 2.5% milk protein.

## DISCUSSION

These investigations began with repeated unsuccessful attempts to form a robust biofilm of *S. marcescens* isolated from soil. Different temperatures, carbon sources, nutrient availability, osmolarity and substrata were tested without effect on biofilm formation. However, one environment in which *S. marcescens* can colonize is the human respiratory system and this provided clues to a possible environmental signal initiating biofilm formation in *Serratia*. Alveolar fluid from human lungs is generally at 5-13% protein (23). This environmental feature of the lung led us to test biofilm formation at several concentrations of protein and identify robust biofilm of *S. marcescens* at 5% milk protein. Moreover, the increase in biofilm productivity was frequently an order of magnitude or greater above that observed in unsupplemented media. Our positive control strain, *P. aeruginosa* PA01, appeared unresponsive to high concentrations of protein in the medium.

These observations were extended to 48 freshwater isolates, four strains from sturgeon eggs (17), two known fish pathogens, three strains isolated from human gut and one from bovine and the results showed biofilm production that was dependent on two variables, species and protein concentration (a total of 74 strains including the *Serratia* isolates). Based on these data *Serratia* isolates from both soil and human gut were highly responsive to 5% exogenous protein, producing 5 to 10 times the amount of biofilm that they made in unsupplemented media. In all cases tested, *Serratia* required concentrations around 5% and failed to respond to lower concentrations (0.1%, 0.2%, 0.4%, 0.5%, 1% & 2%). *Aeromonas* strains were also sensitive to exogenous protein in the same manner, increasing biofilm production, although lower concentrations of protein (0.4% - 2%) would suffice for some strains. The one strain of *F. columnare* reported on herein was particularly responsive to exogenous protein with an optimum at 1% protein and evidence of increased biofilm productivity at as low as 0.1% protein. Additional studies within the *Flavobacterium* and *Chryseobacterium* lineages indicated that all isolates of *F. columnare* tested thus far are responsive to 1% milk protein (Loch & Marsh, unpublished). Biofilm production by *P. aeruginosa* PA01 was unresponsive to high concentrations of protein (1%-5%) and showed growth inhibition but enhanced biofilm production at low concentrations (0.1%, 0.2% & 0.4%). Those strains that appeared unresponsive at high concentrations included freshwater isolates *Kluyvera*, *Erwinia*, nearly all *Pseudomonas* (11 of 12), all *Rahnella aquatilis* isolates, 3 of 4 *Yersinia* isolates and human and bovine *E. coli* isolates.

### Protein as a surface conditioning agent

A number of investigators have reported that soluble protein can serve as a “conditioner” to surfaces that enhance or inhibit the development of biofilm. Frequently serum is used as a “natural” protein-containing solution to condition surfaces (total protein in serum is typically 60-80g/L). For example, Patel et al. (24) showed that initial binding of *S. epidermidis* cells to hydrophobic polyurethanes was suppressed by serum at 2 hours but enhanced when incubated for 24 hrs. The opposite trend was observed for hydrophilic surfaces where serum inhibited biofilm formation. Similarly, Frade et al. (25) found that serum enhanced biofilm productivity of *Candida albicans* on metallic and non-metallic surfaces. Finally, using methodologies most similar to our approach, Kipanga et al. (26) demonstrated that polystyrene microtiter plates (Costar) conditioned with foetal calf serum showed reduced biofilm formation by *C. albicans*. These assays are in general difficult to compare given the diversity of surfaces, strains and complexity of serum. The Patel et al. work used human serum diluted to 20% as the incubation medium whereas Frade et al. and Kipanga et al. used undiluted foetal calf serum only to condition surfaces, but not as the media of incubation. In contrast our experiments used microbiological media grade skim milk protein, autoclaved separately from other media components to eliminate any temperature induced media-protein interactions. Our fully constructed media containing protein up to 5% was used as the incubation media in which biofilm was formed. The observations that different phylogenetic taxa have different optimal protein concentrations for growth, biofilm formation and protein assembled into the biofilm matrix suggest that caution must be used in drawing generalizations regarding the influence of serum (or alveolar fluid) on biofilm formation by any single isolate. The various effective ranges of biofilm enhancement exhibited by different strains in our study suggests that conditioning of the surface was not a relevant factor (our concentrations were beyond saturation levels for polystyrene) but that species dependent sensitivity to protein in the media was driving enhanced biofilm production at various protein concentrations. Direct tests of milk protein as a surface conditioning agent for *Serratia* were negative (data not shown).

### Exogenous protein – a trigger or adjuvant to biofilm formation?

As mentioned above, we were particularly interested in identifying environmental triggers of biofilm formation. While our results with exogenous protein are provocative in this regard, we cannot identify milk protein supplement as a trigger as opposed to an adjuvant in biofilm formation. The experiments described in Figures 5 & 6 clearly indicate that the addition of exogenous protein increased cell concentration within the biofilm matrix (and biofilm biomass as measured with crystal violet) as well as the concentration of matrix protein in *A. salmonicida* and *F. columnare*. *A. salmonicida* was particularly efficient at incorporating protein into the matrix, increasing 80-fold over controls lacking milk protein. Interestingly, the primary strains of this study, *P. aeruginosa*, *A. salmonicida*, *F. columnare* and *S. marcescens*, produce extracellular proteases when grown on R2A or TSA plates with 5% milk protein (data not shown). Other isolates of these strains have a well-documented history of producing extracellular proteases (27–33). Consistent with this was our observations in Fig. 6 that when cultivated in microtiter plates for biofilm production, both *P. aeruginosa* and *F. columnare* reduced the opacity of exogenous protein in the media, indicating that extracellular proteases were actively degrading milk protein under the conditions of our biofilm test. However, *A. salmonicida* showed no such activity in broth but did add an abundance of protein to the biofilm matrix, suggesting that exogenous protein was at least a biofilm adjuvant for *A. salmonicida*. Concluding that exogenous protein is the environmental trigger for *F. columnare* biofilm formation is consistent with the complete absence of detectable pelagic growth in broth supplemented with the milk protein but with concurrent construction of abundant biofilm and incorporation of substantial protein into the matrix. Nonetheless, we do not have direct evidence that exogenous protein is an environmental trigger. Finally, we note that skim milk protein is a common microbiological media additive that is not well defined because of proprietary information claims. The protein concentration range that we employed is not attainable with pure casein.

### Protein, Proteases and Virulence

Some proteases are identified as virulence factors in pathogens including *Serratia* (29) and *Pseudomonas aeruginosa* (30, 33). The simplistic view of these extracellular proteases is that they are foraging for nutrients and clear habitats to occupy as well as impeding host immune responses that are protein based. While extracellular proteases have been linked to biofilm formation in *Enterococcus* (34–36), from our observations it is unclear if extracellular proteases influence the formation of biofilm in *P. aeruginosa, F. columnare*, *S. marcescens* and *A. salmonicida*, under our experimental conditions. With a simple plate assay, we can detect extracellular proteases in these strains but the response to exogenous protein in the production of biofilm is strain specific and *Aeromonas* does not appear to degrade MP in broth when testing for biofilm. Whether or not the proteases generate small peptides that are triggers or adjuvants of biofilm production remains to be determined.

### The host-pathogen evolutionary dance

The analogy of an arms race has been used repeatedly for host-pathogen interactions as they evolve over time (37–40). Within this construct, each actor endeavors to detect the strengths and weaknesses of the other and evolve a strategy that increases the odds of survival, usually at the other’s expense. Biofilm is recognized as a strategic response of bacteria to host defenses in that it protects the inhabitants from antibiotics, host defensins, macrophages and eosinophil networks (5, 10, 41, 42). The studies herein began with *S. marcescens* isolated from soil, a habitat with its own unique set of challenges but one that does not usually include pockets with high concentrations of protein. However, *S. marcescens* is adaptable and can infect both nematodes and humans. In nematodes, infection can initiate in the gut after ingestion (43). Based on the results from our *S. marcescens* strains we would predict that biofilm would be stimulated upon contact with the high protein content of the intestine and the epithelial lining of the nematode (the initial targets for infections caused by *A. salmonicida* and *F. columnare* include the fins, gills and intestinal tract are all sites with elevated protein concentrations). Similarly, in the respiratory system of humans we would predict that *S. marcescens* would form biofilm upon contact with the high protein concentrations of alveolar fluid. With respect to alveolar fluid and infections of the respiratory system, the lung, in contrast to our friendly media with benign milk protein, is designed to be a hostile environment for microbes. The protein content of alveolar fluid is complex and contains many different proteins of which four proteins are abundant, SP-A, SP-B, SP-C, SP-D, and were originally described as hydrophobic (B & C) and hydrophilic (A & D) surfactants that facilitate gas exchange on the mucosal surface (44). These proteins can represent up to 10% of the dry weight of bronchial lavage fluid (45). Of particular interest are SP-A and SP-D, now recognized as collectins, that participate in host defense along with their role as surfactants. Both bind bacterial LPS and in addition, SP-D binds peptidoglycan. These proteins have also been implicated in clearance of pathogens, activation of macrophages, modulation of inflammatory response and regulation of innate immunity functions in the lung (44–46). We posit that the second virulence strategy of *Serratia* (and *Aeromonas*, and *F. columnare*) is the sequestration of proteins from the environment of their host, into the biofilm matrix. This is consistent with the biofilm matrix as a multifunctional extracellular ‘organ’ of a bacterial consortium (47). Incorporation of substantial amounts of SP-A into the biofilm as a structural component would locally reduce its concentration in alveolar fluid and mute the host’s immune response. Targeting SP-A has been previously documented for *P. aeruginosa* (48).

In demonstrating the substantial influence of exogenous protein on biofilm productivity we hope that this stimulates further work on this aspect of biofilm formation. Responses to exogenous protein appeared to be strain specific, suggesting that the different environments in which these strains are colonizing may have a range of exogenous protein concentrations to which cognate strains have adapted. Protein and/or peptides in the concentrations ranges where we have detected enhanced biofilm formation would saturate protein binding sites on the cell surface. Some of these sites, as in the case of *Enteroco*ccus (34, 49, 50), are linked to two-component regulatory systems, hence exogenous protein may be an environmental trigger for biofilm formation.

## Acknowledgements

We thank the Department of Microbiology and Molecular Genetics for partial funding. The remaining funds came through the Water Science program at Michigan State University. This project was conducted almost entirely by undergraduates. The isolation of *Serratia marcescens* strains were performed in an undergraduate laboratory (MMG-302) at MSU. DY, JK, & LB were undergraduates at MSU and responsible for Figures 1, 2, 3, 5 & 6. MMC was a visiting undergraduate scholar from Brazil, supported by Science Without Borders funded by Coordination for the Improvement of Higher Education Personnel and performed experiments for Figures 4 & 9); MS was a visiting undergraduate scholar from Kalamazoo College and performed the initial characterization of the *S. marcescens* isolates.

