## SupplementaryFigs for "Exogenous Protein as an Environmental Stimuli of Biofilm Formation in Select Bacterial Strains"

**Supplemental Files.**
**Figure 1.** Phylogenetic tree of Serratia isolates (RL1-RL16).
**Table 1.** Phylogenetic affiliation of Serratia isolates determined by Ribosomal Database Project Classifier.
**Table 2.** Phylogenetic affiliation of Serratia isolates determined by Ribosomal Database Project Sequence Match.

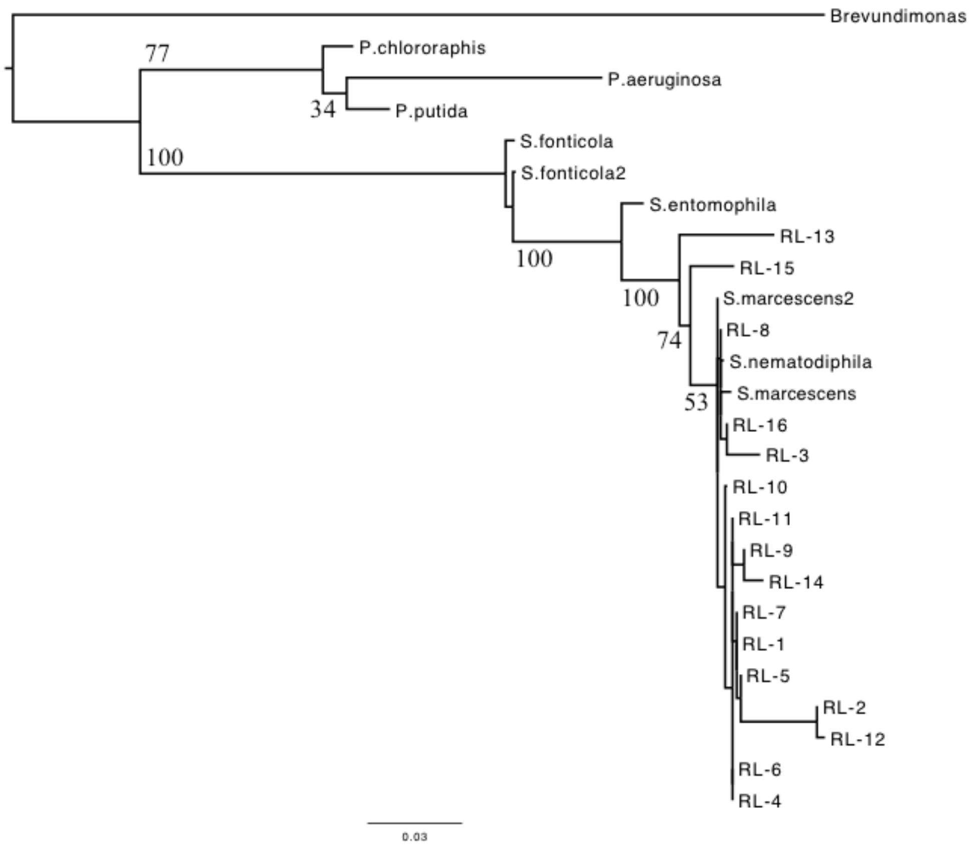


**Figure S1. Maximum Likelihood Tree of 16S rRNA sequences from Serratia isolates (RL-1 – RL-16). Sequences were aligned in SeaView v.4.5.4 (Gouy et al. 2010) and phylogenetically compared using PhyML with GTR model and boot strapped x100. The starting tree was an optimized BioNJ based on 497 positions.**

Gouy, M. Guindon, S. & Gascuel., O. (2010) SeaView version 4 : a multiplatform graphical user interface for sequence alignment and phylogenetic tree building. Molecular Biology and Evolution 27(2):221-224.

**Classifier: RDP Naive Bayesian rRNA Classifier Version 2.11**
**Taxonomical Hierarchy: RDP 16S rRNA training set 16**
**Confidence threshold (for classification to Root ONLY): 80%**
**Symbol +/- indicates predicted sequence orientation**

| | | | | | | | | | | | | |
|---|---|---|---|---|---|---|---|---|---|---|---|---|
| **RL1** | Bacteria | 100% | Proteobacteria | 100% | Gammaproteobacteria | 100% | Enterobacteriales | 100% | Enterobacteriaceae | 100% | Serratia | 100% |
| **RL2** | Bacteria | 100% | Proteobacteria | 100% | Gammaproteobacteria | 100% | Enterobacteriales | 100% | Enterobacteriaceae | 100% | Serratia | 100% |
| **RL3** | Bacteria | 100% | Proteobacteria | 100% | Gammaproteobacteria | 100% | Enterobacteriales | 100% | Enterobacteriaceae | 100% | Serratia | 100% |
| **RL4** | Bacteria | 100% | Proteobacteria | 100% | Gammaproteobacteria | 100% | Enterobacteriales | 100% | Enterobacteriaceae | 100% | Serratia | 100% |
| **RL5** | Bacteria | 100% | Proteobacteria | 100% | Gammaproteobacteria | 100% | Enterobacteriales | 100% | Enterobacteriaceae | 100% | Serratia | 100% |
| **RL6** | Bacteria | 100% | Proteobacteria | 100% | Gammaproteobacteria | 100% | Enterobacteriales | 100% | Enterobacteriaceae | 100% | Serratia | 100% |
| **RL7** | Bacteria | 100% | Proteobacteria | 100% | Gammaproteobacteria | 100% | Enterobacteriales | 100% | Enterobacteriaceae | 100% | Serratia | 100% |
| **RL8** | Bacteria | 100% | Proteobacteria | 100% | Gammaproteobacteria | 100% | Enterobacteriales | 100% | Enterobacteriaceae | 100% | Serratia | 100% |
| **RL9** | Bacteria | 100% | Proteobacteria | 100% | Gammaproteobacteria | 100% | Enterobacteriales | 100% | Enterobacteriaceae | 100% | Serratia | 100% |
| **RL10** | Bacteria | 100% | Proteobacteria | 100% | Gammaproteobacteria | 100% | Enterobacteriales | 100% | Enterobacteriaceae | 100% | Serratia | 100% |
| **RL11** | Bacteria | 100% | Proteobacteria | 100% | Gammaproteobacteria | 100% | Enterobacteriales | 100% | Enterobacteriaceae | 100% | Serratia | 100% |
| **RL12** | Bacteria | 100% | Proteobacteria | 100% | Gammaproteobacteria | 100% | Enterobacteriales | 100% | Enterobacteriaceae | 100% | Serratia | 100% |
| **RL13** | Bacteria | 100% | Proteobacteria | 100% | Gammaproteobacteria | 100% | Enterobacteriales | 100% | Enterobacteriaceae | 100% | Serratia | 98% |
| **RL14** | Bacteria | 100% | Proteobacteria | 100% | Gammaproteobacteria | 100% | Enterobacteriales | 100% | Enterobacteriaceae | 100% | Serratia | 100% |
| **RL15** | Bacteria | 100% | Proteobacteria | 100% | Gammaproteobacteria | 100% | Enterobacteriales | 100% | Enterobacteriaceae | 100% | Serratia | 100% |
| **RL16** | Bacteria | 100% | Proteobacteria | 100% | Gammaproteobacteria | 100% | Enterobacteriales | 100% | Enterobacteriaceae | 100% | Serratia | 100% |

**Table S1. Results from submission of *Serratia* 16S rRNA sequences to RDP Classifier.**

**Seqmatch:version 3**
**RDP Data:release11_5**
**Data Set:both type and non-type strains,**
**:isolates,**
**:near-full-length sequences (>=1200 bases),**
**:good quality sequences**
**Comments:307935 sequences were included in the search. The screening was based on 7-base oligomers.**

| Isolate | S_ab score | unique common oligomers | sequence name | domain | phylum | class | order | family | genus |
|---|---|---|---|---|---|---|---|---|---|
| **RL1** | 0.986 | 1383 | Serratia sp. J21; JN091871 | Bacteria | Proteobacteria | Gammaproteobacteria | Enterobacteriales | Enterobacteriaceae | Serratia |
| **RL2** | 0.958 | 1383 | Serratia sp. J21; JN091871 | Bacteria | Proteobacteria | Gammaproteobacteria | Enterobacteriales | Enterobacteriaceae | Serratia |
| **RL3** | 0.986 | 1367 | Serratia sp. 1135; KC236478 | Bacteria | Proteobacteria | Gammaproteobacteria | Enterobacteriales | Enterobacteriaceae | Serratia |
| **RL4** | 0.982 | 1351 | Serratia marcescens; C2; GU220797 | Bacteria | Proteobacteria | Gammaproteobacteria | Enterobacteriales | Enterobacteriaceae | Serratia |
| **RL5** | 0.975 | 1383 | Serratia sp. J21; JN091871 | Bacteria | Proteobacteria | Gammaproteobacteria | Enterobacteriales | Enterobacteriaceae | Serratia |
| **RL6** | 0.983 | 1351 | Serratia marcescens; C2; GU220797 | Bacteria | Proteobacteria | Gammaproteobacteria | Enterobacteriales | Enterobacteriaceae | Serratia |
| **RL7** | 0.985 | 1364 | Serratia marcescens; VSC-5; HQ130340 | Bacteria | Proteobacteria | Gammaproteobacteria | Enterobacteriales | Enterobacteriaceae | Serratia |
| **RL8** | 1 | 1313 | Serratia marcescens; HO2-A; AJ297950 | Bacteria | Proteobacteria | Gammaproteobacteria | Enterobacteriales | Enterobacteriaceae | Serratia |
| **RL9** | 0.975 | 1351 | Serratia marcescens; C2; GU220797 | Bacteria | Proteobacteria | Gammaproteobacteria | Enterobacteriales | Enterobacteriaceae | Serratia |
| **RL10** | 0.989 | 1351 | Serratia marcescens; C2; GU220797 | Bacteria | Proteobacteria | Gammaproteobacteria | Enterobacteriales | Enterobacteriaceae | Serratia |
| **RL11** | 0.984 | 1351 | Serratia marcescens; C2; GU220797 | Bacteria | Proteobacteria | Gammaproteobacteria | Enterobacteriales | Enterobacteriaceae | Serratia |
| **RL12** | 0.95 | 1364 | Serratia marcescens; VSC-5; HQ130340 | Bacteria | Proteobacteria | Gammaproteobacteria | Enterobacteriales | Enterobacteriaceae | Serratia |
| **RL13** | 0.932 | 1269 | Serratia nematodiphila; KtPC3-5; KF017547 | Bacteria | Proteobacteria | Gammaproteobacteria | Enterobacteriales | Enterobacteriaceae | Serratia |
| **RL14** | 0.957 | 1351 | Serratia marcescens; C2; GU220797 | Bacteria | Proteobacteria | Gammaproteobacteria | Enterobacteriales | Enterobacteriaceae | Serratia |
| **RL15** | 0.934 | 1297 | bacterium 28W333; KC734248 | Bacteria | Proteobacteria | Gammaproteobacteria | Enterobacteriales | Enterobacteriaceae | Serratia |
| **RL16** | 0.99 | 1332 | Serratia marcescens; N1.6; AY514431 | Bacteria | Proteobacteria | Gammaproteobacteria | Enterobacteriales | Enterobacteriaceae | Serratia |

**Table S2. Results from submission of *Serratia* 16S rRNA sequences to RDP SeqMatch Function.**
